# Neural Basis Of Sound-Symbolic Pseudoword-Shape Correspondences

**DOI:** 10.1101/2023.04.14.536865

**Authors:** Deborah A. Barany, Simon Lacey, Kaitlyn L. Matthews, Lynne C. Nygaard, K. Sathian

## Abstract

Non-arbitrary mapping between the sound of a word and its meaning, termed sound symbolism, is commonly studied through crossmodal correspondences between sounds and visual shapes, e.g., auditory pseudowords, like ‘mohloh’ and ‘kehteh’, are matched to rounded and pointed visual shapes, respectively. Here, we used functional magnetic resonance imaging (fMRI) during a crossmodal matching task to investigate the hypotheses that sound symbolism (1) involves language processing; (2) depends on multisensory integration; (3) reflects embodiment of speech in hand movements. These hypotheses lead to corresponding neuroanatomical predictions of crossmodal congruency effects in (1) the language network; (2) areas mediating multisensory processing, including visual and auditory cortex; (3) regions responsible for sensorimotor control of the hand and mouth. Right-handed participants (*n* = 22) encountered audiovisual stimuli comprising a simultaneously presented visual shape (rounded or pointed) and an auditory pseudoword (‘mohloh’ or ‘kehteh’) and indicated via a right-hand keypress whether the stimuli matched or not. Reaction times were faster for congruent than incongruent stimuli. Univariate analysis showed that activity was greater for the congruent compared to the incongruent condition in the left primary and association auditory cortex, and left anterior fusiform/parahippocampal gyri. Multivoxel pattern analysis revealed higher classification accuracy for the audiovisual stimuli when congruent than when incongruent, in the pars opercularis of the left inferior frontal (Broca’s area), the left supramarginal, and the right mid-occipital gyri. These findings, considered in relation to the neuroanatomical predictions, support the first two hypotheses and suggest that sound symbolism involves both language processing and multisensory integration.

**HIGHLIGHTS:**

- fMRI investigation of sound-symbolic correspondences between auditory pseudowords and visual shapes
- Faster reaction times for congruent than incongruent audiovisual stimuli
- Greater activation in auditory and visual cortices for congruent stimuli
- Higher classification accuracy for congruent stimuli in language and visual areas
- Sound symbolism involves language processing and multisensory integration

## INTRODUCTION

Sound symbolism refers to the idea that the sounds of a word resemble its meaning (Nuckolls, 1999). This idea dates back to Plato’s Cratylus dialog (Ademollo, 2011), but has receded in modern times in favor of the mainstream view that sound-meaning associations are essentially arbitrary (e.g. Pinker, 1999). Recently, however, there has been a resurgence of interest in the phenomenon, fueled by the discovery of crossmodal correspondences between the sounds of words and (non-auditory) perceptual dimensions. For instance, auditory pseudowords such as ‘takete’ or ‘kiki’ are reliably matched with pointed visual shapes, whereas ‘maluma’ or ‘bouba’ are matched with rounded shapes (Köhler, 1929, 1947; Ramachandran & Hubbard, 2001). Further, the perceptual ratings of auditory pseudowords as rounded or pointed relate not only to phonological features (e.g., McCormick et al., 2015; Fort et al., 2015; Cuskley et al., 2017), but also to their acoustic properties, including spectrotemporal parameters of the pseudoword sounds and parameters of vocal roughness associated with sound production (Lacey et al., 2020).

Prior functional magnetic resonance imaging (fMRI) studies measuring the blood oxygenation level-dependent (BOLD) signal investigated the neural basis of sound symbolism using a variety of tasks across different semantic domains, without much consistency of findings. Revill et al. (2014) asked native English speakers to listen to words drawn from multiple foreign languages and match their meanings with English antonym pairs spanning domains of motion, size, and shape. In this study, greater activity in the left intraparietal sulcus for sound-symbolic compared to non-sound-symbolic words was considered to reflect crossmodal correspondences between object properties and word sounds, based on the recruitment of a nearby area during audiovisual synesthetic experiences (Neufeld et al., 2012). Visually presented Japanese mimetic words for motion and shape, when they matched visual images of actions or shapes, evoked activity in the right posterior superior temporal sulcus; this was construed as indicating conceptual integration of language and non-language sounds (Kanero et al., 2014). Judging the size of visual objects during presentation of sound symbolically incongruent, relative to congruent, pseudowords led to greater activity in the right anterior superior temporal gyrus and the left posterior middle temporal gyrus: the authors interpreted their data in terms of reliance on phonological and semantic processes (Itagaki et al., 2019). Congruent tactile hard/soft stimuli and visually presented Japanese words that were sound-symbolic for hardness/softness, relative to incongruent stimulus pairs, led to activity in areas involved in tactually assessing softness: the anterior insula and medial superior frontal gyrus bilaterally (Kitada et al., 2021). These regions also distinguished between congruent and incongruent stimuli in a multivoxel pattern analysis (MVPA) (Kitada et al., 2021), though the reported analyses did not explore whether individual stimuli could be distinguished. Collectively, these studies drew inferences about the relevant neural processes, but they were not designed to evaluate alternative explanations.

In the sound-symbolic shape domain, two previous fMRI studies presented participants with concurrent auditory pseudowords and visual shapes. Peiffer-Smadja & Cohen (2019) reported greater activity for incongruent than congruent audiovisual pairs in the dorsolateral prefrontal cortex bilaterally in a task where participants were asked to detect occasional target stimuli unrelated to the pseudoword-shape correspondence of interest, possibly reflecting greater effort during processing of the incongruent stimuli. In a prior study from our group, McCormick et al. (2022) found that incongruent stimuli elicited stronger BOLD responses than congruent stimuli in frontoparietal regions when participants attended to the auditory input; these findings were thought to be consistent with changes in phonological processing and/or multisensory attention as a function of sound-symbolic congruency (McCormick et al., 2022). Although this study was conducted to arbitrate between Spence’s (2011) accounts of crossmodal correspondences as originating in audiovisual statistical regularities, a domain-general magnitude system, or similarity of the pseudowords to real words, clear evidence did not accrue for or against any of these accounts.

In the present fMRI study, we investigated the neural correlates of sound symbolism for shape when participants were asked to explicitly assess the sound-symbolic audiovisual pseudoword-shape correspondence. Effects favoring congruent over incongruent stimuli were not observed in the two studies cited above, which used implicit matching of auditory pseudowords and visual shape (Peiffer-Smadja & Cohen, 2019; McCormick et al., 2022), but did occur during explicit matching of visually presented words to visual or tactile stimuli (Kanero et al., 2014; Kitada et al., 2021). Thus, we reasoned that an explicit judgment task, by focusing attention on whether the stimuli matched or not, would be more likely to elicit congruency effects. In addition to examining univariate BOLD effects, we also performed MVPA. We predicted that classification accuracy for specific pseudoword-shape combinations would be higher in congruent than incongruent trials and that regions distinguishing between congruent and incongruent stimuli would be distinct from those differentiating between specific stimulus pairs.

Our experimental approach allowed us to examine three hypotheses, which are not necessarily mutually exclusive, regarding the processes involved in perceiving sound-symbolic audiovisual pseudoword-shape correspondences:

**Hypothesis 1** draws on a large body of psychophysical work implicating sound symbolism in language structure and function, e.g. the ubiquity of sound-symbolic associations across multiple languages (Blasi et al., 2016), the higher frequency of rounded (pointed) phonemes in words signifying rounded (pointed) objects (Sidhu et al., 2021), the facilitation of word learning (Nielsen & Dingemanse, 2021; Nygaard et al., 2009) and associative sound-shape memory (Sonier et al., 2020) by sound symbolism, the human ability to create sound-symbolic words for actions seen in locomotion videos (Saji et al., 2019), the capability of English speakers to infer Pokemon types based on sound-symbolic Japanese names (Kawahara et al., 2021), and the relative advantage for onomatopoeic words in aphasia (Meteyard et al., 2015). Such findings fit with Hypothesis 1, that *sound symbolism recruits language processing*, and lead to the neuroanatomical prediction that congruency effects would be found in classical language areas. While this hypothesis draws some support from prior neuroimaging studies (Kanero et al., 2014; Itagaki et al., 2019; see above), these studies did not evaluate alternative possibilities.

**Hypothesis 2** proposes that *multisensory integrative processes underlie sound-symbolic audiovisual correspondences*, due to statistical associations between the auditory and visual stimuli (Spence, 2011; Fort & Schwartz, 2022). Due to these associations, congruent audiovisual pairings lead to multisensory integration whereas incongruent pairings do not (Spence, 2011). As reviewed above, this hypothesis is consistent with the findings of Revill et al. (2014), although this study did not seek to assess other hypotheses. It predicts congruency effects not only in areas classically thought to mediate multisensory processing, but also in areas traditionally regarded as unisensory, i.e. visual and auditory cortex, given the wealth of evidence for widely distributed multisensory interactions in the neocortex (Ghazanfar & Schroeder, 2006).

**Hypothesis 3** is rooted in the idea of acoustic similarities between pseudowords that are matched to rounded or pointed shapes and the sounds of manual actions used to draw such shapes with a sound-producing pen (Margiotoudi & Pulvermüller, 2020). Studies in the size domain suggest a different kind of relationship between articulatory and manual movements: foreign-language words or pseudowords that sound symbolically imply small or large size are associated with a precision or power grip, respectively (Vainio & Vainio, 2021, 2022; Vainio et al., 2023). These studies gave rise to Hypothesis 3, that *sound symbolism is based on embodiment of speech in hand actions*. This hypothesis, not previously tested in neuroimaging work, makes the neuroanatomical prediction that brain regions mediating motor control and related somatosensory processing for the hand and mouth are candidate loci for congruency effects.

## METHODS

### Participants

Twenty-four people took part in this study (14 female, 10 male; mean age = 22.6 years, SD = 3.8 years). All participants were right-handed based on the validated subset of the Edinburgh handedness inventory (Raczkowski et al., 1974) and reported normal hearing and normal, or corrected-to-normal, vision. All participants gave written informed consent and were compensated for their time. All procedures were approved by the Emory University Institutional Review Board and complied with the Declaration of Helsinki. Participants performed three scanning sessions; in the first two, auditory pseudowords and visual shapes were presented in separate sessions (which are the subject of a separate report), and the third comprised the audiovisual task reported here. Two participants with poor behavioral performance (see *Multisensory fMRI task*) were excluded from further analyses.

### Stimuli

We created sound-symbolic audiovisual stimuli, each consisting of an auditory pseudoword and a two-dimensional visual shape. Two auditory pseudowords (‘mohloh’ and ‘kehteh’) and two shapes (pointed and rounded) were chosen from large stimulus sets (McCormick et al., 2015, see below; McCormick, unpublished data, see below: Figure 1A). The pseudowords and shapes chosen were near the extremes of independent rounded and pointed dimensions based on empirical ratings for 537 pseudowords (McCormick et al., 2015) and 90 visual shapes (McCormick, unpublished data). These independent rating scales were converted to a single scale on which ‘most rounded’ = 1 and ‘most pointed’ = 7; ratings for the pseudowords ranged from 2.3 – 5.8 (median 4.2) and for the shapes from 1.6 – 6.7 (median 4.4). ‘Mohloh’ was the 12^th^ most rounded pseudoword, rated 2.8, but we used the 20th most pointed pseudoword, ‘kehteh’ (rated 5.4), in order to get the closest match in duration. We used the 10^th^ most rounded visual shape, rated 2.3, and the 10^th^ most pointed visual shape, rated 6.4, to match the number of protuberances (five) between the shapes. A more detailed analysis of the pseudowords and shapes can be found in Lacey et al. (2020) and the stimuli themselves are available at https://osf.io/ekpgh/.

**Figure 1.**
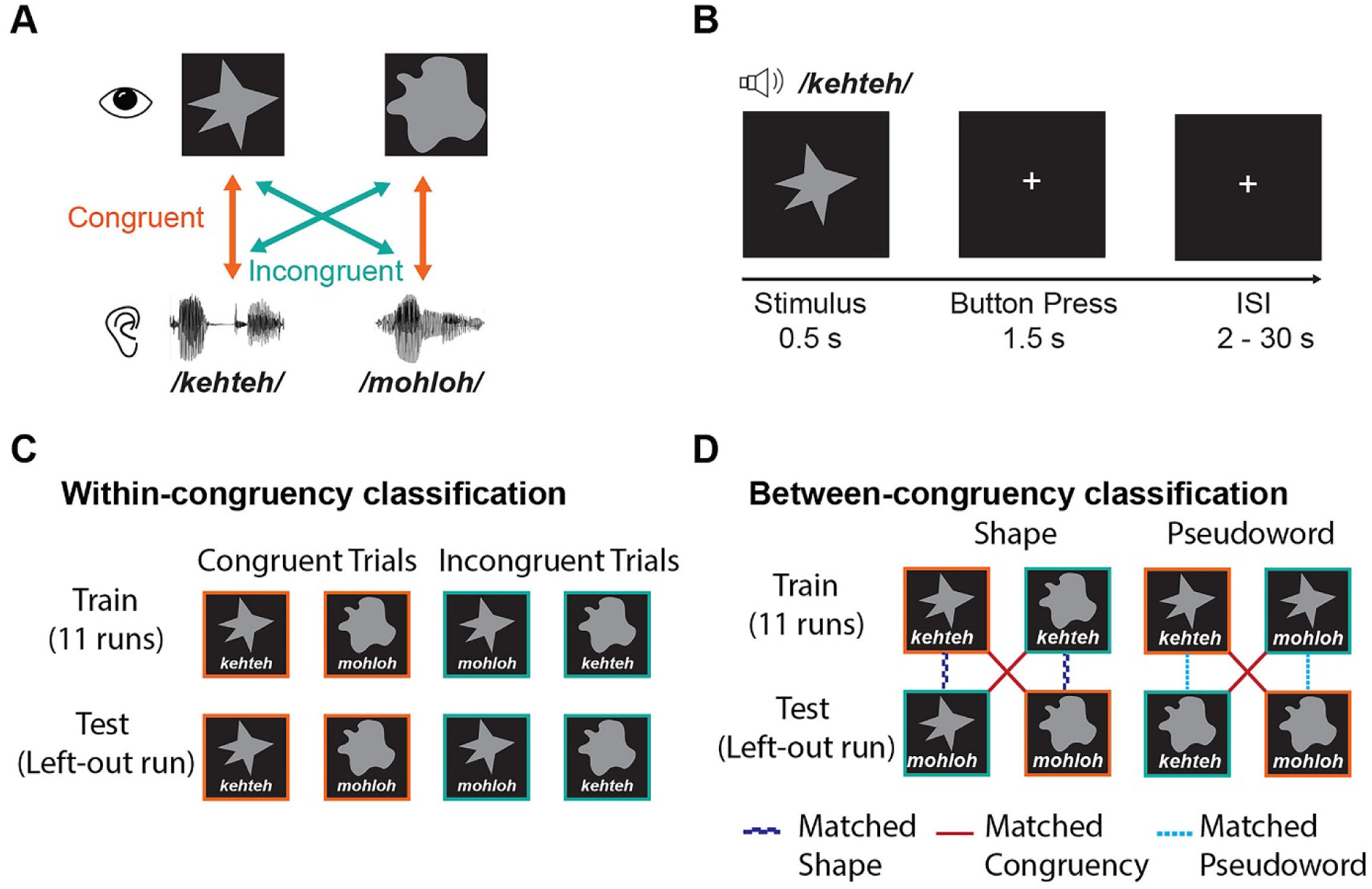
Task design and classification analysis scheme. (A) There were four possible stimuli, reflecting unique combinations of two visual shapes, pointed or rounded, and two auditory pseudowords, ‘kehteh’ or ‘mohloh’. (B) Pseudoword-shape pairs were presented simultaneously and participants pressed a response button indicating whether the pair were a ‘match’ (i.e., congruent) or a ‘mismatch’ (i.e., incongruent). (C) In the within-congruency classification analysis, leave-one-run-out cross-validation was used to test for patterns of brain activity discriminating the two congruent conditions or the two incongruent conditions. (D) in the between-congruency classification analyses, classifiers were trained to distinguish between the two conditions in which the visual shapes differed but the auditory pseudoword was the same (e.g., ‘kehteh’/pointed shape vs. ‘kehteh’/rounded shape). The classifier was then tested on the two remaining conditions containing the other pseudoword (e.g., ‘mohloh’/pointed shape vs. ‘mohloh’/rounded shape). Similarly, we trained classifiers to distinguish between auditory pseudowords holding the shape constant (e.g., ‘kehteh’/pointed shape vs. ‘mohloh’/pointed shape) and then tested them on the other shape (e.g., ‘kehteh’/rounded shape vs. ‘mohloh’/rounded shape). This approach differentiates whether patterns of brain activity were specific to features of the visual/auditory stimulus, or to the congruency/incongruency of the condition.

The pseudowords were digitally recorded in a female voice using Audacity v2.0.1 (Audacity Team, 2012), with a SHURE 5115D microphone and an EMU 0202 USB external sound card, at a 44.1 kHz sampling rate. The recordings were then processed in Sound Studio (Felt Tip Inc., NY), using standard tools and default settings, edited into separate files, amplitude-normalized, and down-sampled to a 22.05 kHz sampling rate (standard for speech). Stimulus duration was 563 ms for ‘mohloh’ and 546 ms for ‘kehteh’, resulting in a difference in duration of 17ms or approximately 3%. Because sound segments in language naturally differ in duration, these differences were retained. The visual shapes were solid gray shapes on a black background, each subtending approximately 1° of visual angle and presented at the center of the screen for 500ms. Duration of visual shape presentation was not varied to match the accompanying pseudowords, to avoid having the shape duration vary as a function of congruency and thus introduce a confound.

Stimuli were presented concurrently in audiovisual pairs (Figure 1A) that were either congruent (‘mohloh’/rounded shape or ‘kehteh’/pointed shape) or incongruent (‘kehteh’/rounded shape or ‘mohloh’/pointed shape) with respect to the crossmodal pseudoword-shape (sound-symbolic) correspondence. Stimuli were delivered via MATLAB 2014b (The MathWorks Inc., Natick, MA) to synchronize delivery with fMRI acquisition and also record responses and reaction times (RTs). A mirror angled over the head coil enabled participants to see the visual stimuli projected onto a screen placed in the rear magnet aperture. Auditory stimuli were presented via scanner-compatible Sensimetrics S14 headphones (Sensimetrics Corporation, Gloucester MA).

### Multisensory fMRI task

We used a jittered event-related design in which participants performed 12 runs of the audiovisual task. Each run consisted of 22 congruent trials and 22 incongruent trials, each contained within the TR of 2 s (see Image acquisition, below: Figure 1B), giving a trial sample size, across runs, of 132 for each of the four unique audiovisual pairs (see Figure 1A). This number of trials per condition is over 3 times the upper end of the typical range (10-40) and is large enough to permit condition-specific modeling of responses (Chen et al., 2022). Inter-trial intervals (ITIs), or null periods, ranged from 2 s to 30 s. We used optseq2 (https://surfer.nmr.mgh.harvard.edu/optseq/; Dale, 1999; Dale et al., 1999) to schedule trials and ITIs, in order to optimize the estimation of the hemodynamic response. Each run began and ended with a rest period of 10 s for a total run duration of 326 s. Pseudoword-shape pairs were presented in a fixed pseudorandom order within each run; the 12 run orders were counterbalanced across participants. Participants attended to each audiovisual stimulus pair and judged whether the pair was a match or a mismatch, pressing one of two buttons on a hand-held response box to indicate their response. Participants were free to make their own judgments of match/mismatch; they were not informed of our interest in studying sound symbolism. The right index and middle fingers were used to indicate match/mismatch responses, counterbalanced across subjects.

All participants, except two, judged the ‘match’ condition to be congruent pairs of stimuli and the ‘mismatch’ condition to be incongruent pairs of stimuli. These two participants responded with the opposite than expected mapping for at least one of the pseudoword-shape pairs; their data were excluded from analysis.

### Image acquisition

MR scans were performed on a 3 Tesla Siemens Trio TIM whole body scanner (Siemens Medical Solutions, Malvern, PA), using a 32-channel head coil. T2*-weighted functional images were acquired using a single-shot, gradient-recalled, echoplanar imaging (EPI) sequence for BOLD contrast. For functional scans, 34 axial slices of 3.1 mm thickness were acquired using the following parameters: repetition time (TR) 2000 ms, echo time (TE) 30 ms, field of view (FOV) 200 mm, flip angle (FA) 90°, in-plane resolution 3.125×3.125 mm, and in-plane matrix 64×64. High-resolution 3D T1-weighted (T1w) anatomic images were acquired using an MPRAGE sequence (TR 2300 ms, TE 3.9ms, inversion time 1100 ms, FA 8°) comprising 176 sagittal slices of 1 mm thickness (FOV 256 mm, in-plane resolution 1×1 mm, in-plane matrix 256×256). Once magnetic stabilization was achieved in each run, the scanner triggered the computer running MATLAB and the Psychophysics Toolbox extensions (Brainard, 1997; Pelli, 1997; Kleiner et al, 2007) so that the experiment was synchronized with scan acquisition.

### Image preprocessing

Results included in this manuscript come from preprocessing performed using *fMRIPrep* version 20.0.6, (Esteban et al., 2019; Esteban et al., 2020; RRID:SCR_016216), which is based on *Nipype* 1.4.2 (Gorgolewski et al., 2011; RRID:SCR_002502). Many internal operations of *fMRIPrep* use *Nilearn* 0.6.2 (Abraham et al., 2014, RRID:SCR_001362], mostly within the functional processing workflow. For more details of the pipeline, see the section corresponding to workflows in *fMRIPrep*’s documentation (https://fmriprep.readthedocs.io/en/latest/workflows.html). The T1w image was corrected for intensity non-uniformity (INU) with N4BiasFieldCorrection (Tustison et al., 2010), distributed with ANTs 2.2.0 (Avants et al., 2008; RRID:SCR_004757), and used as the T1w-reference throughout the workflow. The T1w-reference was then skull-stripped with a *Nipype* implementation of the antsBrainExtraction.sh workflow (from ANTs), using OASIS30ANTs as the target template. Brain tissue segmentation of cerebrospinal fluid (CSF), white matter (WM) and gray matter (GM) was performed on the brain-extracted T1w using ‘fast’ (Zhang et al., 2001; FSL 6.0.3:b862cdd5, RRID:SCR_002823).

For each of the 12 BOLD runs per subject, the following preprocessing was performed.

First, a reference volume and its skull-stripped version were generated using a customized implementation of *fMRIPrep*. Susceptibility distortion correction (SDC) was omitted. The BOLD reference was then co-registered to the T1w reference using ‘flirt’ (FSL 6.0.3:b862cdd5, Jenkinson & Smith, 2001) with the boundary-based registration (Greve & Fischl, 2009) cost function. Co-registration was configured with nine degrees of freedom to account for distortions remaining in the BOLD reference. Head-motion parameters with respect to the BOLD reference (transformation matrices, and six corresponding rotation and translation parameters) were estimated before spatiotemporal filtering using mcflirt (FSL 6.0.3:b862cdd5, Jenkinson et al., 2002). BOLD runs were slice-time corrected using 3dTshift from AFNI 20190316 (Cox, 1996, RRID:SCR_005927). The BOLD time-series (including slice-timing correction when applied) were resampled onto their original, native space by applying the transforms to correct for head motion. These resampled BOLD time-series will be referred to as preprocessed BOLD in original space, or just preprocessed BOLD.

Several confounding time-series were calculated based on the preprocessed BOLD: framewise displacement (FD), the derivative of the root mean squared variance over voxels (DVARS), and three region-wise global signals. FD and DVARS were calculated for each functional run, using their implementations in *Nipype* (following the definitions by Power et al., 2014). The three global signals were extracted within the CSF, the WM, and the whole-brain masks. Additionally, a set of physiological regressors was extracted to allow for component-based noise correction (*CompCor*, Behzadi et al., 2007). Principal components were estimated after high-pass filtering the preprocessed BOLD time-series (using a discrete cosine filter with 128 s cut-off) for the two *CompCor* variants: temporal (tCompCor) and anatomical (aCompCor). tCompCor components were then calculated from the top 5% variable voxels within a mask covering the subcortical regions. This subcortical mask was obtained by heavily eroding the brain mask, which ensures it does not include cortical GM regions. For aCompCor, components were calculated within the intersection of the aforementioned mask and the union of CSF and WM masks calculated in T1w space, after their projection to the native space of each functional run (using the inverse BOLD-to-T1w transformation). Components were also calculated separately within the WM and CSF masks. For each CompCor decomposition, the *k* components with the largest singular values were retained, such that the retained components’ time series were sufficient to explain 50 percent of variance across the nuisance mask (CSF, WM, combined, or temporal). The remaining components were dropped from consideration.

The head-motion estimates calculated in the correction step were also placed within the corresponding confounds file. The confound time series derived from head motion estimates and global signals were expanded with the inclusion of temporal derivatives and quadratic terms for each (Satterthwaite et al., 2013). Frames that exceeded a threshold of 0.5 mm FD or 1.5 standardised DVARS were annotated as motion outliers. All resamplings were performed with a single interpolation step by composing all the pertinent transformations (i.e. head-motion transform matrices, susceptibility distortion correction when available, and co-registrations to anatomical and output spaces). Gridded (volumetric) resamplings were performed using antsApplyTransforms (ANTs), configured with Lanczos interpolation to minimize the smoothing effects of other kernels (Lanczos, 1964). Non-gridded (surface) resamplings were performed using ‘mri_vol2surf’ from FreeSurfer.

### fMRI data analysis

Statistical analyses of fMRI data were conducted using SPM12 (Wellcome Trust Center for Neuroimaging, London, UK) as implemented in *Nipype*. We used a standard General Linear Model (GLM) to estimate separate regressors (betas) for each of the four conditions in each run. Stimulus onsets were modeled as a boxcar function with duration equal to that trial’s reaction time (Grinband et al., 2008) and convolved with the canonical hemodynamic response function. The six translation and rotation head-motion parameter estimates were included as additional regressors of no interest.

A univariate analysis was conducted by first smoothing the preprocessed BOLD signal in native space with an 8 mm full-width half-maximum (FWHM) isotropic Gaussian kernel prior to the first-level analysis. We then performed a linear contrast to evaluate the difference in BOLD activity between the congruent (‘mohloh’/rounded shape or ‘kehteh’/pointed shape) and incongruent (‘kehteh’/rounded shape or ‘mohloh’/pointed shape) conditions. The resulting individual contrast maps were normalized to a standard MNI brain with 1×1×1 mm voxels and then submitted to a second-level analysis of group data with subject treated as a random factor (so that the degrees of freedom equal n-1, i.e., 21), followed by pairwise contrasts. An explicit mask based on a combined gray matter and white matter mask from segmentation of the T1w images was applied at the group level to isolate within-brain activations. Correction for multiple comparisons (*p* <0.05) was achieved using the topological FDR correction method, using whole brain activation clusters with cluster-forming threshold of *p* < 0.001 and FDR cluster-level correction of *q* < 0.05 (Chumbley et al., 2010).

Searchlight multivariate pattern analyses (MVPA) were performed using The Decoding Toolbox (TDT) in SPM (Hebart et al., 2015). The unsmoothed (Misaki et al., 2013) GLM run-wise beta estimates for each of the four conditions were first z-scored voxelwise and within each run to factor out univariate effects, avoid spurious correlations between mean estimates across runs, and improve classification accuracy (Lee & Kable, 2018; Stehr et al., 2023). We implemented two types of cross-validated, leave-one-run-out searchlight decoding analyses using spherical searchlights with a 4-voxel radius and an L2-norm regularized support vector machine classifier with *C* = 1 from the LIBSVM package (Chang & Lin, 2011). For the *within-congruency classification*, the classifier was trained to discriminate between either the two congruent stimulus pairs (i.e., ‘kehteh’/pointed shape vs. ‘mohloh’/rounded shape) or the two incongruent ones (i.e., ‘mohloh’/pointed shape vs. ‘kehteh’/rounded shape) separately (Figure 1C). To directly test whether any region showed significantly better decoding when the stimuli were congruent relative to when they were incongruent (or vice versa), we contrasted the individual classification accuracy maps for the congruent conditions with those for the incongruent conditions.

To test for specific patterns of brain activity related to congruency independent of sensory characteristics, we performed a *between-congruency classification* (Kaplan et al., 2015; Man et al., 2015). To decode the visual shapes, classifiers were trained to distinguish between two of the four conditions in which the visual shapes were different but the auditory pseudoword was the same (e.g., ‘kehteh’/pointed shape vs. ‘kehteh’/rounded shape: Figure 1D-Shape). The classifier was then tested on the two remaining conditions containing the other pseudoword (e.g., ‘mohloh’/pointed shape vs. ‘mohloh’/rounded shape: Figure 1D-Shape). Regions that represent visual properties should show decoding of visual shape independent of the accompanying auditory pseudoword (e.g., similar patterns of activity for ‘kehteh’/rounded shape and ‘mohloh’/rounded shape). In contrast, in regions where responses are driven by the congruency of crossmodal correspondences (or the associated motor responses), we should be able to decode the congruency/incongruency of the stimuli (e.g., similar patterns of activity for ‘kehteh’/pointed shape and ‘mohloh’/rounded shape). Similarly, to decode congruency/incongruency across auditory features, we trained classifiers to distinguish between auditory pseudowords when the accompanying visual shape was held constant (e.g., ‘kehteh’/pointed shape vs. ‘mohloh’/pointed shape) and then tested on the two remaining conditions containing the other shape (e.g., ‘kehteh’/rounded shape vs. ‘mohloh’/rounded shape: Figure 1D-Pseudoword). As before, we should be able to differentiate regions where auditory features drive responses from regions that represent crossmodal congruency independent of the pseudoword presented.

For both the *within-* and *between-congruency* analyses, the individual classification accuracy searchlight maps were smoothed with a 4 mm full-width half-maximum (FWHM) isotropic Gaussian kernel and normalized to MNI space before second-level analysis of group data with subject as a random factor. All analyses included a combined GM/WM explicit mask to exclude non-brain voxels. For the *within-congruency* analysis comparing differences in classification accuracy for congruent > incongruent (incongruent > congruent), only voxels exhibiting significant above-chance classification of the congruent (incongruent) conditions at the group level were included. The group-level maps were corrected for multiple comparisons using the topological FDR correction method with cluster-forming threshold of *p* < 0.001 and FDR cluster-level correction of *q* < 0.05 (Chumbley et al., 2010).

All activations were localized with respect to 3D cortical anatomy with the AAL (Tzourio-Mazoyer et al., 2002; Rolls et al., 2015) and AICHA (Joliot et al., 2015) atlases; differences between the two were resolved by reference to an MRI atlas (Cho et al., 2010). Activation maps are overlaid on a group-averaged structural image normalized to MNI space (1×1×1 mm voxels).

## RESULTS

### Behavioral data

Table 1 shows the mean accuracy and RTs for each of the four conditions. Accuracy was not significantly different for congruent vs. incongruent trials (*t*_21_ = 0.261; *p* = 0.797; *d* = 0.056). RTs were faster when the pseudoword and shape were sound symbolically congruent compared to when they were incongruent (*t*_21_ = 5.374; *p* < 0.001; *d* = 1.15).

**Table 1.** Mean (standard error) accuracy and reaction times.

| Stimulus pairs | Accuracy (%) | Reaction Time (ms) |
| --- | --- | --- |
| **Congruent** |  |  |
| 'mohloh'/rounded shape | 95.21 (0.75) | 898 (33) |
| 'kehteh'/pointed shape | 95.73 (0.79) | 832 (31) |
| **Incongruent** |  |  |
| 'mohloh'/pointed shape | 96.35 (0.99) | 958 (31) |
| 'kehteh'/rounded shape | 94.11 (1.76) | 938 (33) |

### fMRI univariate analyses

The contrast of congruent > incongruent conditions showed higher activation in the right, and near the left, caudate nucleus, as well as in the left anterior fusiform/parahippocampal gyri, right mid-cingulate gyrus/corpus callosum, left mid-superior temporal gyrus, and left postcentral gyrus. The reverse contrast (incongruent > congruent) showed greater activation in the left precentral sulcus, left supplementary motor area (SMA), bilateral anterior insula, and right pons. These activations were cluster-corrected for multiple comparisons (cluster-forming threshold *p* < 0.001; FDR cluster-level *q* < 0.05) (Figure 2 and Table 2).

**Figure 2.**
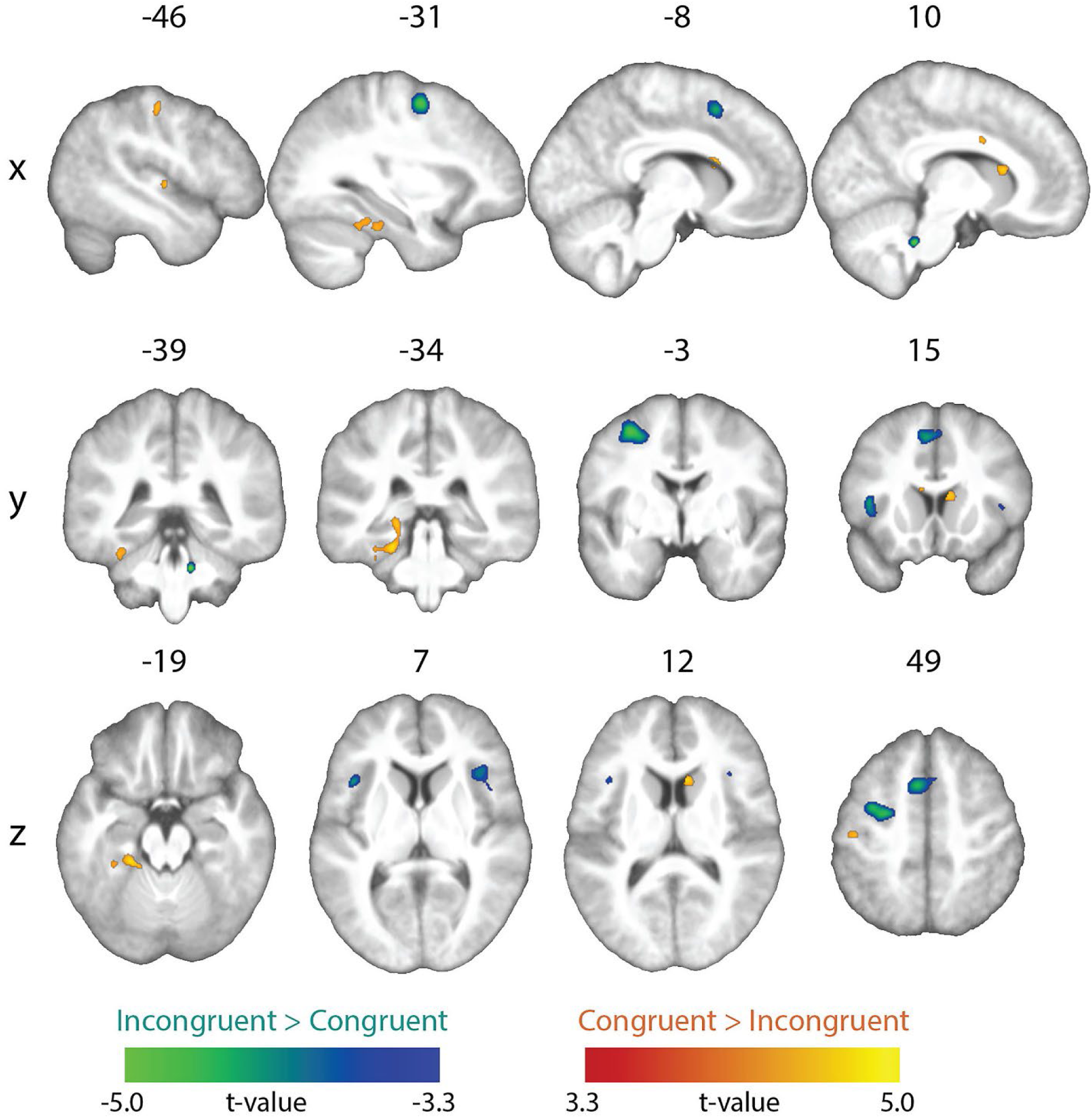
Univariate results. Whole-brain map showing significant effects from the incongruent vs. congruent contrasts (topological FDR-corrected; activation clusters with cluster-forming threshold of *p* < 0.001 and FDR cluster-level correction of *q* < 0.05).

**Table 2.** Brain regions showing univariate activation differences for congruent vs. incongruent stimuli (whole-brain topological FDR-corrected; activation clusters with cluster-forming threshold of *p* < 0.001 and FDR cluster-level correction of *q* < 0.05). Coordinates of peak local maxima are shown for each cluster.

| Anatomical Location | Peak MNI coordinates |  |  | Cluster Level |  |
| --- | --- | --- | --- | --- | --- |
| | x | y | z | mm<sup>3</sup> | $P_{FDR-corr}$ |
| **Congruent > incongruent** |  |  |  |  |  |
| L postcentral gyrus | -45 | -19 | 49 | 163 | < 0.001 |
| L mid-superior temporal gyrus | -46 | -13 | 2 | 49 | 0.016 |
| L anterior fusiform/parahippocampal gyrus | -21 | -34 | -19 | 929 | < 0.001 |
| L caudate nucleus | -8 | 14 | 17 | 71 | 0.013 |
| R mid-cingulate gyrus/corpus callosum | 10 | 3 | 31 | 54 | 0.013 |
| R caudate nucleus | 9 | 16 | 12 | 153 | < 0.001 |
| **Incongruent > congruent** |  |  |  |  |  |
| L precentral sulcus | -31 | -3 | 52 | 1456 | < 0.001 |
| L supplementary motor area | -6 | 13 | 49 | 984 | < 0.001 |
| L anterior insula | -41 | 15 | 8 | 434 | < 0.001 |
| R anterior insula | 38 | 21 | 7 | 576 | < 0.001 |
| R pons | 11 | -39 | -32 | 136 | < 0.001 |

### fMRI searchlight MVPA

#### Within-congruency classification

In separate decoding analyses of the congruent and incongruent conditions, a whole-brain searchlight analysis showed robust above-chance classification accuracy in visual and auditory cortices, reflecting distinct patterns for unique visual shapes and auditory pseudowords, respectively (Figure 3; Table 3). To test where decoding of shape and pseudowords was better for congruent than incongruent trials, we contrasted the individual classification accuracy maps for the congruent condition with those for the incongruent condition. This contrast was restricted to those voxels that exhibited above-chance classification in the congruent condition. The resulting difference map (cluster-corrected for multiple comparisons with cluster-forming threshold *p* < 0.001; FDR cluster-level *q* < 0.05) revealed significantly higher above-chance classification in the congruent, compared to the incongruent, conditions in the pars opercularis of the left inferior frontal gyrus (IFG) (Broca’s area), the left supramarginal gyrus (SMG), and the right mid-occipital gyrus (MOG), (Figure 4A-C respectively; Table 4). Classifier performance differences between congruent and incongruent conditions in Broca’s area and the left SMG resulted from above-chance (∼60% accuracy) classification for the congruent, but not the incongruent, conditions (Figure 4A & B respectively). Classification accuracies at the peak coordinate in the right MOG cluster were above chance for both congruent and incongruent conditions, but significantly higher for the congruent condition, as shown by the bar graphs in Figure 4C. Note that the right MOG cluster was part of a large swath of occipital (early visual) cortex with significant classifier accuracy in the congruent condition. No region showed significantly higher above-chance classification of the incongruent conditions after correction for multiple comparisons.

**Figure 3.**
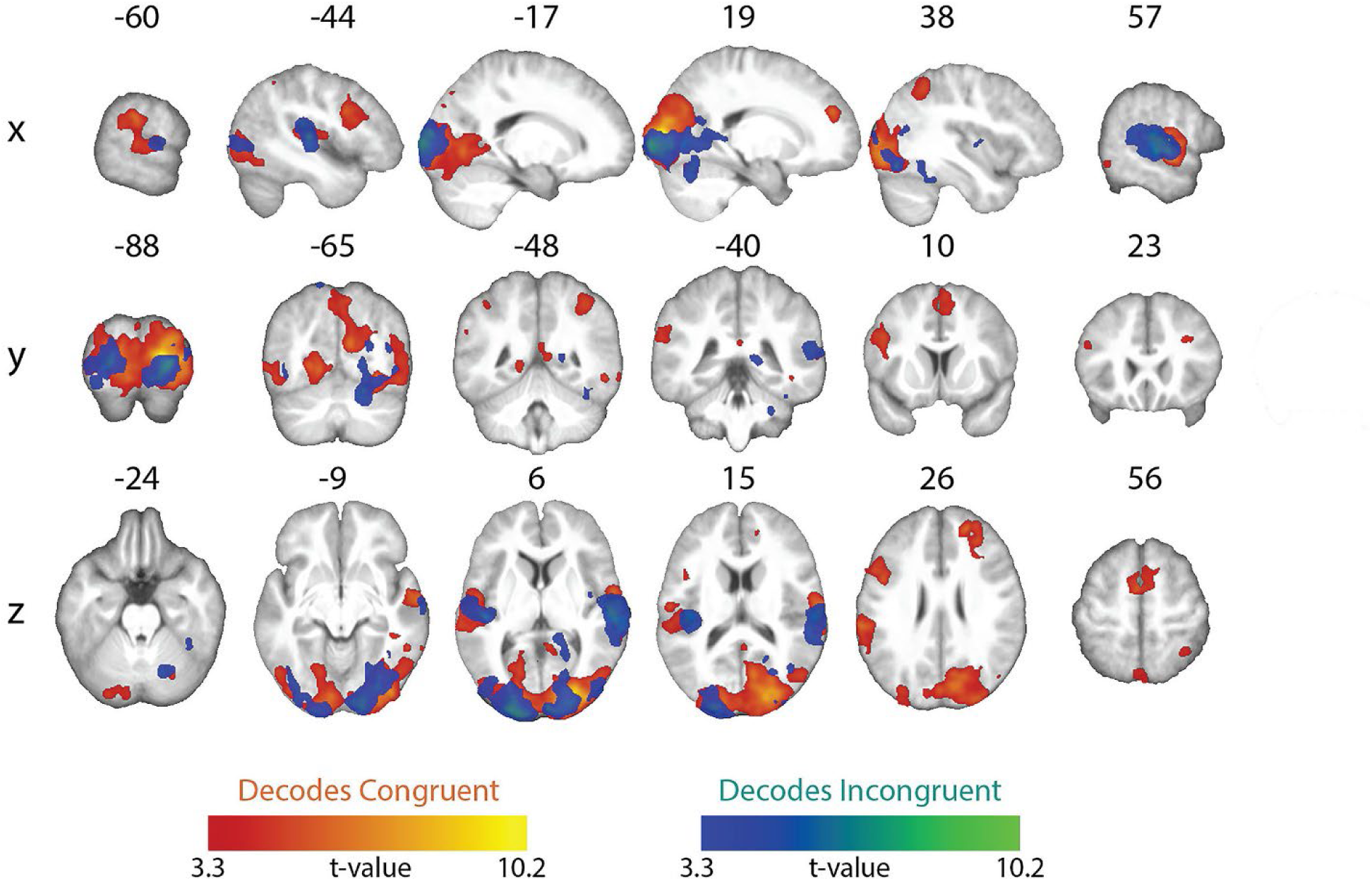
Whole-brain searchlight maps showing significant above-chance within-congruency classification of the two congruent conditions (congruent decoding) and the two incongruent conditions (incongruent decoding). All maps whole-brain topological FDR-corrected; activation clusters with cluster-forming threshold of *p* < 0.001 and FDR cluster-level correction of *q* < 0.05).

**Figure 4.**
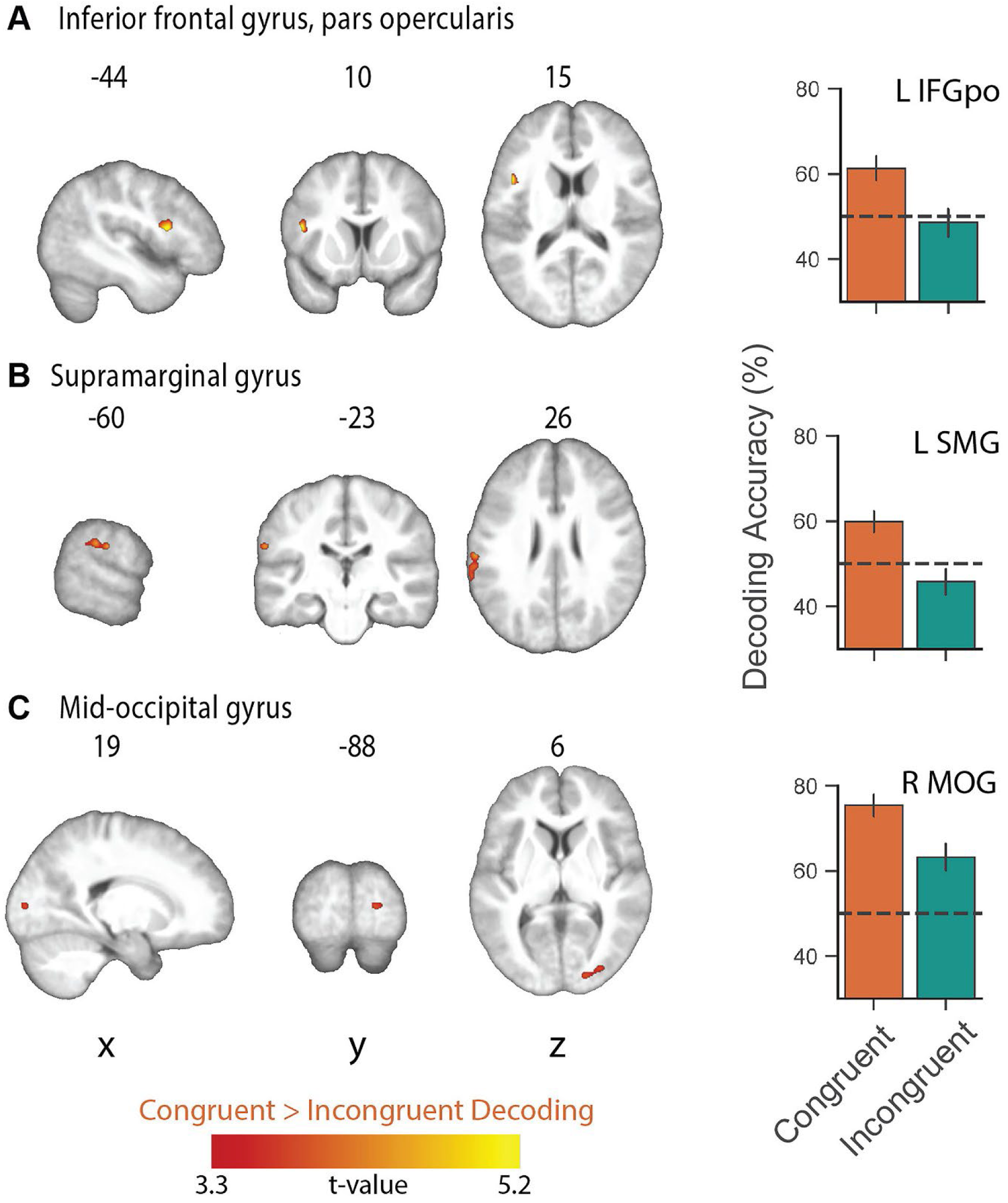
Whole-brain searchlight map showing significantly higher classification of the congruent conditions relative to the incongruent conditions and mean decoding accuracy in (A) left inferior frontal gyrus, pars opercularis (Broca’s area: L IFGpo), (B) left supramarginal gyrus (L SMG), and (C) right mid-occipital gyrus (MOG). Mean decoding accuracy was extracted from the peak voxel coordinates for each of the significant clusters in A-C. No region showed significantly higher above-chance classification of the incongruent conditions. All maps whole-brain topological FDR-corrected; activation clusters with cluster-forming threshold of *p* < 0.001 and FDR cluster-level correction of *q* < 0.05); error bars show ± standard error of the mean.

**Table 3.** Brain regions showing above-chance classification of visual and auditory stimuli in congruent and incongruent conditions (whole-brain topological FDR-corrected; activation clusters with cluster-forming threshold of *p* < 0.001 and FDR cluster-level correction of *q* < 0.05). Coordinates of peak local maxima are shown for each cluster.

| Anatomical Location | Peak MNI coordinates |  |  | Cluster Level |  |
| --- | --- | --- | --- | --- | --- |
| | x | y | z | mm<sup>3</sup> | $P_{FDR-corr}$ |
| **Congruent decoding** |  |  |  |  |  |
| L supplementary motor area | -2 | 0 | 56 | 4171 | < 0.001 |
| L inferior frontal gyrus, pars opercularis | -45 | 6 | 28 | 3263 | < 0.001 |
| L intraparietal sulcus | -41 | -50 | 49 | 228 | 0.006 |
| L supramarginal gyrus | -62 | -30 | 26 | 13328 | < 0.001 |
| R superior frontal gyrus | 20 | 46 | 25 | 3704 | < 0.001 |
| R intraparietal sulcus | 38 | -49 | 48 | 2096 | < 0.001 |
| R precuneus | 6 | -47 | 6 | 583 | 0.006 |
| R superior temporal gyrus | 57 | -12 | -5 | 10332 | < 0.001 |
| R middle temporal gyrus | 64 | -48 | -9 | 122 | 0.042 |
| R inferior temporal gyrus | 53 | -47 | -10 | 162 | 0.020 |
| R inferior temporal gyrus | 41 | -38 | -9 | 152 | 0.023 |
| R calcarine sulcus | 19 | -87 | 7 | 1080150 | < 0.001 |
| **Incongruent decoding** |  |  |  |  |  |
| L superior temporal gyrus | -42 | -27 | 18 | 7938 | < 0.001 |
| L precuneus | -13 | -65 | 66 | 251 | 0.004 |
| L middle occipital gyrus | -17 | -100 | 4 | 21732 | < 0.001 |
| R superior temporal gyrus | 59 | -26 | 11 | 15147 | < 0.001 |
| R fusiform gyrus | 38 | -42 | -24 | 450 | < 0.001 |
| R middle occipital gyrus | 40 | -63 | 15 | 259 | 0.004 |
| R calcarine sulcus | 16 | -98 | -1 | 26326 | < 0.001 |
| R cerebellum | 26 | -40 | -35 | 150 | 0.026 |

**Table 4.**
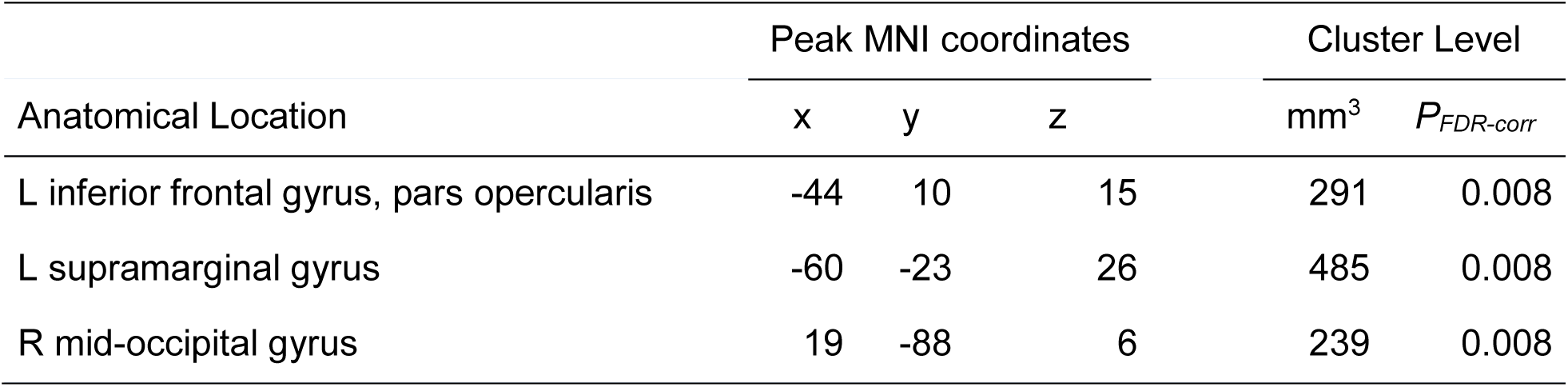
Brain regions showing higher above-chance classification of visual and auditory stimuli in the congruent conditions vs. the incongruent conditions (whole-brain topological FDR-corrected; activation clusters with cluster-forming threshold of *p* < 0.001 and FDR cluster-level correction of *q* < 0.05). Coordinates of peak local maxima are shown for each cluster. Note: no region showed greater incongruent > congruent decoding.

| Anatomical Location | Peak MNI coordinates |  |  | Cluster Level |  |
| --- | --- | --- | --- | --- | --- |
| | x | y | z | mm<sup>3</sup> | $P_{FDR-corr}$ |
| L inferior frontal gyrus, pars opercularis | -44 | 10 | 15 | 291 | 0.008 |
| L supramarginal gyrus | -60 | -23 | 26 | 485 | 0.008 |
| R mid-occipital gyrus | 19 | -88 | 6 | 239 | 0.008 |

#### Between-congruency classification

A whole-brain searchlight analysis revealed distinct regions for processing of the visual shapes and auditory pseudowords, and for processing their crossmodal correspondences (Figure 5, Tables 5 and 6). When classifiers were trained to distinguish the two visual shapes, significant above-chance decoding of visual shape was restricted to visual cortical regions with peak local maxima in the lingual gyrus and calcarine sulcus, whereas decoding of crossmodal congruency (i.e. congruent vs. incongruent) was significantly above chance in multiple regions, including left precentral gyrus, bilateral superior frontal gyri, right pallidum, right postcentral gyrus, left hippocampus, right superior parietal lobule, right middle frontal gyrus, left posterior cingulate gyrus, and right inferior frontal gyrus (Figure 5A, Table 5). A similar pattern of results was observed for classifiers trained to distinguish between the auditory pseudowords: only bilateral auditory cortex (with peak local maxima in the superior temporal sulcus/gyrus) displayed significant above-chance pseudoword decoding, whereas there was significant above-chance decoding of crossmodal congruency/incongruency in multiple bilateral parietofrontal and temporal regions, and on the right in the parahippocampal gyrus, cerebellum, thalamus, and anterior insula (Figure 5B, Table 6). Notably, congruency decoding in parietofrontal regions was more prominent in the left hemisphere (contralateral to the index and middle fingers used to pressing the buttons indicating match or mismatch responses).

**Figure 5.**
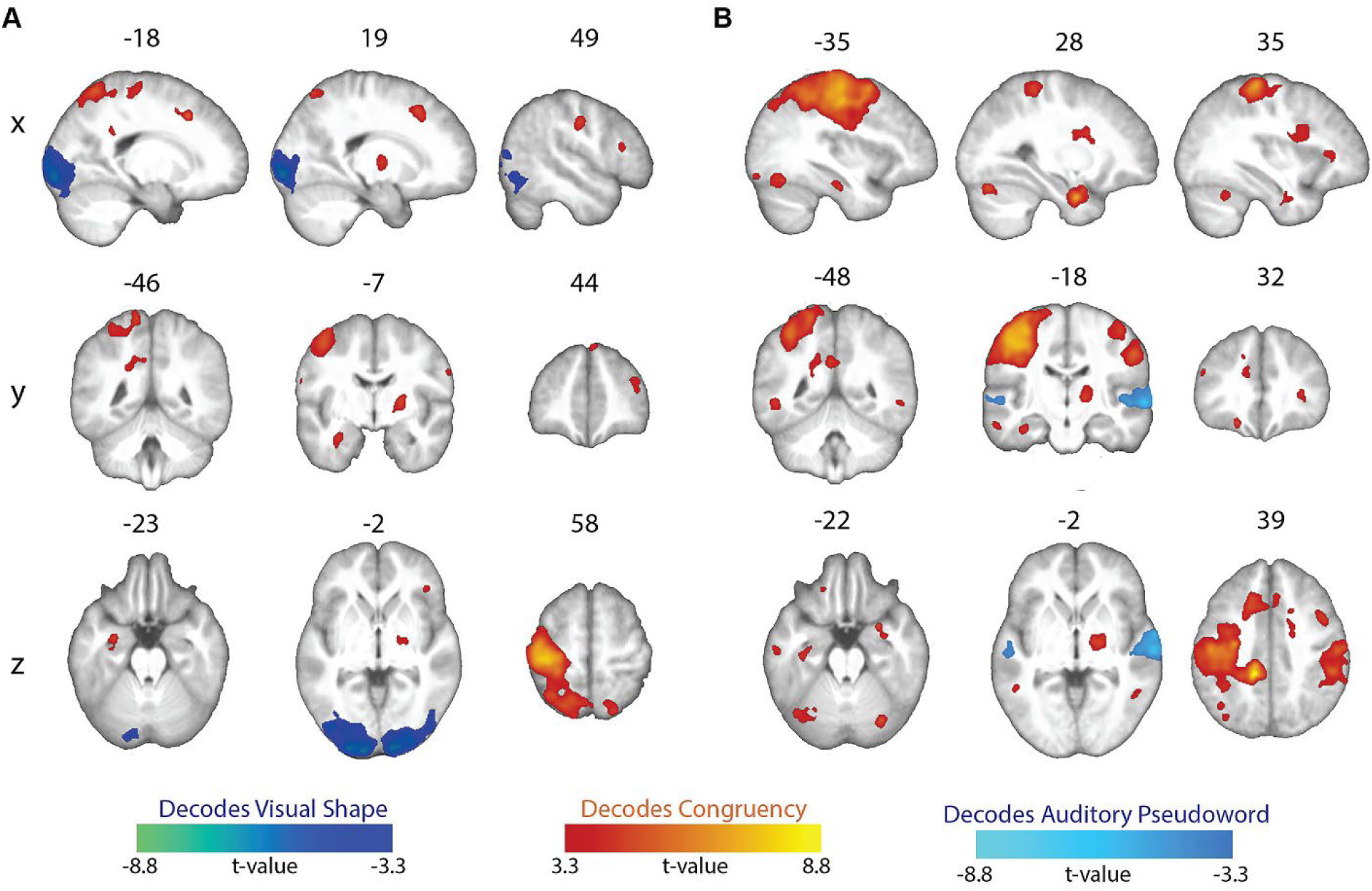
Between-congruency classification. (A) Whole-brain searchlight maps showing significant above-chance decoding of either the visual shape or the congruency of the stimuli. (B) Whole-brain searchlight maps showing significant above-chance decoding of either the auditory pseudoword or the congruency of the stimuli. All maps whole-brain topological FDR-corrected; activation clusters with cluster-forming threshold of *p* < 0.001 and FDR cluster-level correction of *q* < 0.05).

**Table 5.** Brain regions showing above-chance decoding of visual shape and congruency (whole-brain topological FDR-corrected; activation clusters with cluster-forming threshold of *p* < 0.001 and FDR cluster-level correction of *q* < 0.05). Coordinates of peak local maxima are shown for each cluster.

| Anatomical Location | Peak MNI coordinates |  |  | Cluster Level |  |
| --- | --- | --- | --- | --- | --- |
| | x | y | z | mm<sup>3</sup> | $P_{FDR-corr}$ |
| **Shape decoding** |  |  |  |  |  |
| L calcarine sulcus | -11 | -99 | -4 | 20279 | < 0.001 |
| R lingual gyrus | 16 | -93 | -6 | 19354 | < 0.001 |
| **Congruency decoding** |  |  |  |  |  |
| L superior frontal gyrus | -17 | 17 | 41 | 752 | < 0.001 |
| L precentral gyrus | -35 | -25 | 60 | 42895 | < 0.001 |
| L posterior cingulate gyrus | -19 | -46 | 28 | 1050 | < 0.001 |
| L hippocampus | -29 | -6 | -23 | 706 | < 0.001 |
| R superior frontal gyrus | 5 | 45 | 50 | 151 | 0.023 |
| R superior frontal gyrus | 19 | 20 | 44 | 1476 | < 0.001 |
| R middle frontal gyrus | 38 | 41 | 23 | 832 | < 0.001 |
| R inferior frontal gyrus, pars triangularis | 49 | 22 | 15 | 280 | 0.007 |
| R inferior frontal gyrus, pars orbitalis | 41 | 32 | -2 | 121 | 0.043 |
| R postcentral gyrus | 49 | -13 | 35 | 1813 | < 0.001 |
| R superior parietal lobule | 20 | -67 | 58 | 627 | < 0.001 |
| R pallidum | 23 | -7 | 6 | 928 | < 0.001 |

**Table 6.**
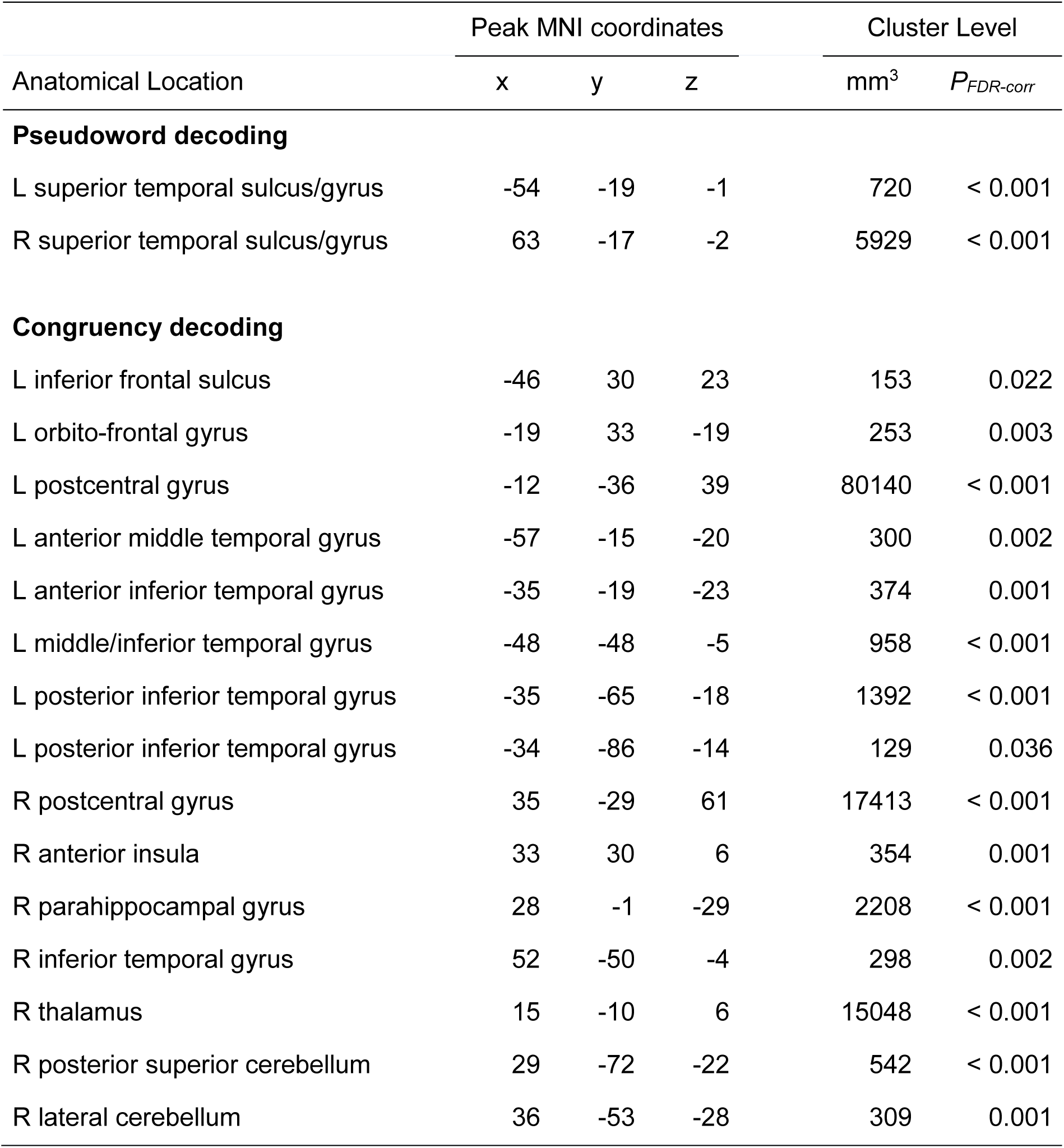
Brain regions showing above-chance decoding of auditory pseudowords and congruency (whole-brain topological FDR-corrected; activation clusters with cluster-forming threshold of *p* < 0.001 and FDR cluster-level correction of *q* < 0.05). Coordinates of peak local maxima are shown for each cluster.

## DISCUSSION

Previous fMRI studies of the sound-symbolic correspondence between auditory pseudowords and visual shapes (McCormick et al., 2022; Peiffer-Smadja & Cohen, 2019) were limited to univariate analyses of task conditions involving implicit registration of the correspondence. The present study is the first fMRI investigation of this psychophysically well-studied sound-symbolic correspondence using an explicit audiovisual matching task and incorporating MVPA to supplement conventional univariate analyses. MVPA offers insights into much finer-grained information processing than is possible with conventional univariate analyses, and has not previously been applied to the sound-symbolic pseudoword-shape correspondence, or to decode particular stimulus combinations in neuroimaging studies of sound symbolism. Our study used an event-related design and had a slightly higher participant sample size (n=22) than the prior studies, both of which employed block designs (McCormick et al., 2022: n=19; Peiffer-Smadja & Cohen, 2019: n=18). Importantly, our study also had a high number of trials (132) per unique condition, allowing for reliable condition-level modeling of responses and making a major contribution to statistical power (Chen et al., 2022). These features of experimental design and analysis enable novel insights into the neural basis of sound symbolism.

We found some neuroanatomical support for each of the hypotheses outlined in the Introduction, based on concordance between the areas showing univariate and multivariate congruency effects and the anatomical predictions derived from the hypotheses. As predicted *a priori*, MVPA classification accuracy was higher for congruent than incongruent conditions in certain areas, but the reverse was not true. **Hypothesis 1**, proposing that sound symbolism is associated with language processing, and thus predicting congruency effects in classical language areas, was supported by MVPA revealing superior classification performance of the bisensory stimulus pairs when congruent, compared to when they were incongruent, in regions of the classical language network: the pars opercularis of the left IFG, i.e., Broca’s area, and the left SMG (Figure 4A & B respectively, Table 4). Our findings also support **Hypothesis 2**, suggesting that sound symbolism depends on multisensory integration and therefore recruits sensory cortical processes for congruent stimuli, in classical multisensory areas as well as areas regarded conventionally as unisensory. Evidence for this hypothesis comes from the MVPA producing higher classifier accuracy for congruent relative to incongruent conditions in early visual cortex (Figure 4C, Table 4), and higher univariate magnitudes of the BOLD signal for the congruent than the incongruent condition in primary and association auditory cortex and in a part of higher-order visual cortex (Figure 2, Table 2). There was limited support for **Hypothesis 3**, the notion that sound symbolism arises from embodiment of speech in manual actions, and the consequent prediction of congruency effects in motor and somatosensory regions related to the hand and mouth. The only data potentially consistent with this idea were the univariate demonstration of greater BOLD signal favoring the congruent over the incongruent condition in the hand area of primary somatosensory cortex (postcentral gyrus) (Figure 2, Table 2). However, the absence of the predicted concomitant activity in regions mediating motor control weakens support for this hypothesis.

Earlier univariate fMRI studies of the pseudoword-shape correspondence (McCormick et al., 2022; Peiffer-Smadja & Cohen, 2019) did not find regions that were preferentially active for congruent relative to incongruent stimuli, perhaps owing to reliance on implicit matching tasks. Here, our explicit matching task revealed univariate activations for congruent compared to incongruent stimuli, as Kitada et al. (2021) found in touch-related regions when participants judged whether tactile stimuli were matched or mismatched with Japanese mimetic words related to tactile hardness and softness.

A valuable feature of our MVPA was testing whether activity patterns were related to low-level sensory features or to processing congruency. We observed above-chance between-congruency classification of the two visual stimuli in visual cortical areas, regardless of which auditory stimulus was paired with them. Similar above-chance classification of the two pseudoword stimuli was observed in auditory cortical areas, regardless of the accompanying visual stimulus. These findings (Figure 5, Table 5 & 6) are not surprising, but provide direct evidence that such neural processing of low-level visual/auditory features is distinct from the processing of congruency discussed below. Classification of stimulus pairs as congruent or incongruent, regardless of the particular stimuli, was above chance in multiple brain regions.

Among these regions, frontoparietal areas in the left hemisphere were prominent, perhaps attributable to decoding of finger movements (Shen et al., 2014), since each participant always used one finger to indicate that the audiovisual stimuli matched (congruent), and another finger to indicate the stimuli were mismatched (incongruent). However, loci classifying congruent vs. incongruent extended beyond the motor network and may relate to variations in attentional demand or decision processes for distinguishing congruent from incongruent stimuli. Future work is needed to isolate regions sensitive to processing congruency independent of the motor response.

### Sound symbolism involves language processing

As predicted by **Hypothesis 1**, MVPA classifier accuracy on the bisensory stimulus pairs was higher for congruent pairs in Broca’s area and the left SMG, both regions of the classical language network (Figure 4A & B, Table 4). In these areas, the classifier yielded above-chance performance for congruent, but not incongruent stimuli. These findings support the idea that sound symbolism is associated with language processing.

Although Broca’s area is involved in syntactic processing (Fedorenko & Blank, 2020; Kemmerer, 2022; Maran et al., 2022), a number of studies also implicate it in audiovisual associations relevant to language: For instance, high-gamma power in Broca’s area was enhanced by congruent audiovisual speech, with a progressive increase in coherence between auditory cortex and Broca’s area over the duration of the stimulus, whereas coherence between these regions decreased over time for incongruent audiovisual speech (Lange et al., 2013).

Another study found that anodal transcranial direct current stimulation over Broca’s area facilitated acquisition of novel associations between pseudowords and unfamiliar visual objects (Perikova et al., 2022). An fMRI study using MVPA found that viewing of congruent lip movements facilitated classification of phonemes in Broca’s area, with a larger effect for phonemes distinguished by voicing (voiced vs. unvoiced consonants) than by place of articulation (Zhang and Du, 2022). Our finding of higher classification performance for congruent, compared to incongruent, pseudoword-shape pairs in Broca’s area resonates with these earlier studies and provides evidence that processing of audiovisual congruency related to sound symbolism is indeed associated with language (Blasi et al., 2016; Kawahara et al., 2021; Meteyard et al., 2015; Nielsen & Dingemanse, 2021; Nygaard et al., 2009; Saji et al., 2019; Sidhu et al., 2021; Sonier et al., 2020).

Similar reasoning may apply to the left SMG, where again classification of the bisensory stimulus pairs was better when they were congruent rather than incongruent (Figure 4B, Table 4). The SMG has been implicated in language processing, particularly phonological processes (Hartwigsen et al., 2010; Oberhuber et al., 2016). Reinforcing this line of thought, Zhang and Du (2022) found that viewing congruent lip movements facilitated classification of phonemes in the left SMG in addition to Broca’s area; however, in the SMG, classifier performance was significant for place of articulation but not voicing. The left SMG also demonstrates beta power suppression for incongruent audiovisual speech (Lange et al., 2013). Curiously, a number of studies have demonstrated stronger univariate activation for incongruent than congruent stimuli in both Broca’s area (Ojanen et al., 2005; Pekkola et al., 2006; Szycik et al., 2009) and the left SMG (Bernstein et al., 2009; Xie et al., 2019; McCormick et al, 2022), which may reflect greater effort associated with processing of incongruent audiovisual stimuli related to speech. Though we did not observe increased univariate activation in Broca’s area or left SMG in the present study, our MVPA findings of superior classifier performance for congruent compared to incongruent stimuli suggest that the neuronal populations recruited in these areas are more distinct for congruent than incongruent audiovisual stimuli, independent of overall activation.

### Involvement of multisensory integration in sound symbolism

In the right MOG, in early visual cortex, the MVPA classifier performed above chance for both congruent and incongruent stimulus pairs, with better performance for congruent pairings (Figures 3 & 4C, Tables 3 & 4). This is consistent with multisensory convergence with a preference for congruency, indicating a role for multisensory integration in sound symbolism, in accord with **Hypothesis 2**.

Some of the univariate activations for congruent relative to incongruent stimuli in the present study were in relevant sensory cortical areas (Figure 2, Table 2): One focus was in auditory cortex in the superior temporal gyrus, including primary auditory cortex (Moerel et al., 2014) and neighboring auditory association cortex (Malinowska et al., 2017). Another was in a higher-order region of visual cortex, in the anterior fusiform gyrus (see Kliger & Yovel, 2020), extending into the parahippocampal gyrus, which has been implicated in evaluating audiovisual congruency (Diaconescu et al., 2011). We also observed preferential bilateral activation near the caudate nucleus for congruent relative to incongruent audiovisual stimuli. Although the mechanism underlying this caudate nucleus activation is unclear, a similar preference for audiovisual congruency in the caudate nuclei has been found previously (Stevenson et al., 2009; McNorgan et al, 2015). Thus, these activations may reflect multisensory convergence (e.g., Schroeder et al., 2003), with audiovisual congruency leading to stronger activity in these regions, in keeping with **Hypothesis 2**.

### Embodiment of sound symbolism in manual actions

The left somatosensory cortical focus in the postcentral gyrus displayed higher univariate activity for congruent relative to incongruent stimuli (Figure 2, Table 2). Its anatomical location was in the hand representation (Martuzzi et al., 2014), medial to both the face representation (Moulton et al., 2009) and the part of somatosensory cortex activated during listening to and repeating speech (Darainy et al., 2019). In keeping with **Hypothesis 3**, activity at this focus could be due to postulated embodiment of the roundedness/pointedness of auditory pseudowords in manual actions, given similarities between the sounds of shape-signifying pseudowords and of hand actions employed in drawing corresponding shapes (Margiotoudi & Pulvermüller, 2020). However, if so, it is curious that the primary motor cortex or areas mediating motor planning were not recruited. Thus, support for **Hypothesis 3** is weak, at least in the shape domain of sound symbolism explored here. It remains possible that this hypothesis may be more applicable to the size domain of sound symbolism, based on the line of work indicating the association of pseudowords that sound symbolically imply small or large size with a precision or power grip, respectively (Vainio & Vainio, 2021, 2022; Vainio et al., 2023); this may be worth testing in future neuroimaging studies.

### Processing of incongruent audiovisual stimuli

Greater activity for incongruent, compared to congruent, bimodal stimuli was found in cortex of the left precentral sulcus (premotor cortex) and the left SMA (Figure 2, Table 2), both regions involved in motor planning. Recruitment of these regions cannot be attributed solely to RT differences in the motor response since RT duration for each trial was explicitly modeled in the GLM (Grinband et al., 2008). While these activations could be construed to support **Hypothesis 3**, this interpretation would be at odds with that offered in the preceding paragraph, and the preponderance of evidence in the present study for both univariate and multivariate effects favoring the congruent condition. An alternative possibility is that greater activation in motor areas for incongruent trials could be due to the reprogramming of a preplanned response to congruent stimuli (Hartwigsen et al., 2012; Pellegrino et al., 2018).

Finally, we found higher activity for incongruent relative to congruent stimuli in the dorsal region of the anterior insula bilaterally (Figure 2, Table 2). In a meta-analytic study, this region was functionally connected with the anterior cingulate cortex and dorsolateral prefrontal cortex, and was associated with higher cognitive tasks and executive control (Chang et al., 2013).

Anterior insula activation may also reflect a domain-general processing of surprise (Loued-Khenissi et al., 2020). These activations may hence be related to those reported in earlier studies using implicit sound-symbolic matching, where bimodally incongruent stimuli elicited stronger activation than congruent stimuli in prefrontal cortical areas and the intraparietal sulcus, postulated to indicate neural processes related to attention or effort (McCormick et al., 2022; Peiffer-Smadja & Cohen, 2019).

Additional research is required to test the inferences made regarding processing of incongruent stimuli, since these were not associated with well-defined predictions.

### Limitations

We used an explicit judgment task to uncover neural responses related to the processing of congruency that were not observed in previous studies relying on implicit matching (Peiffer-Smadja & Cohen, 2019; McCormick et al., 2022). A caveat to this approach is that neural activity associated with the motor response required for the explicit judgment may confound detection of condition-specific effects. To address this, we adopted a standard approach used for both univariate analysis and MVPA to account for RT differences across congruent and incongruent conditions (Grinband et al., 2008; Woolgar et al., 2015). However, this method assumes that the BOLD signal scales with RT during the task, which may mask congruency-related activity that is relatively constant across trials (Mumford et al., 2023). Further, we restricted our stimuli to two pseudowords and two shapes that were distinctly associated with “roundness” or “pointedness” based on empirical ratings, but were otherwise well-matched on stimulus parameters (Lacey et al., 2020). This allowed us to clearly define pseudoword-shape correspondences for simple explicit judgments and to maximize our trial sample size for statistical efficiency (Chen et al., 2022), but may limit the generalizability of our findings.

Although our interpretations of the fMRI findings rely on reverse inference, they do so in the context of a particular task and anatomical priors (Hutzler, 2014), based in hypotheses that arise from the psychophysical literature. Future replications of the present work, use of condition-rich designs, probing of hypothesized neural processes, and extension to other sound-symbolic domains is necessary to more fully understand how crossmodal congruency representations may vary across a continuum of stimuli.

### Conclusions

This fMRI study provided evidence for multiple neural processes underlying sound symbolism. Most interestingly, the present study offers support for a relationship of sound symbolism to the neural processes mediating language and multisensory integration. Other relevant operations include lower-level auditory and visual cortical processing of the stimuli, somatic sensorimotor processing, and more general cognitive operations mediating attention or decisions. Additional work is needed to disentangle the contributions of these various processes to sound symbolism and to more fully understand the role of sound symbolism in language.

## ACKNOWLEDGMENTS

This work was supported by grants to KS and LCN from the National Eye Institute at the NIH (R01 EY025978) and the Emory University Research Council. Support to KS from the Veterans Administration is also acknowledged. Neuroimaging data were collected at the Facility for Education and Research in Neuroscience (FERN) at Emory University. We thank Kate Pirog Revill for assistance with scanning and Sara List for data collection.

## DATA AND CODE AVAILABILITY

Data are available on request to the corresponding author and on submission of a completed data sharing agreement, project outline, and agreement on any co-authorship requirements.

